# Interplay between antibiotic efficacy and drug-induced lysis underlies enhanced biofilm formation at subinhibitory drug concentrations

**DOI:** 10.1101/163733

**Authors:** Wen Yu, Kelsey Hallinen, Kevin B. Wood

**Affiliations:** Department of Physics, University of Michigan, Ann Arbor, Michigan 48109, USA; Department of Biophysics, University of Michigan, Ann Arbor, Michigan 48109, USA

## Abstract

Subinhibitory concentrations of antibiotics have been shown to enhance biofilm formation in multiple bacterial species. While antibiotic exposure has been associated with modulated expression in many biofilm-related genes, the mechanisms of drug-induced biofilm formation remain a focus of ongoing research efforts and may vary significantly across species. In this work, we investigate antibiotic-induced biofilm formation in *E. faecalis*, a leading cause of nosocomial infections. We show that biofilm formation is enhanced by subinhibitory concentrations of cell wall synthesis inhibitors, but not by inhibitors of protein, DNA, folic acid, or RNA synthesis. Furthermore, enhanced biofilm is associated with increased cell lysis, an increase in extracellular DNA (eDNA), and an increase in the density of living cells in the biofilm. In addition, we observe similar enhancement of biofilm formation when cells are treated with non-antibiotic surfactants that induce cell lysis. These findings suggest that antibiotic-induced biofilm formation is governed by a trade-off between drug toxicity and the beneficial effects of cell lysis. To understand this trade-off, we developed a simple mathematical model that predicts changes to antibiotic-induced biofilm formation due to external perturbations, and we verify these predictions experimentally. Specifically, we demonstrate that perturbations that reduce eDNA (DNase treatment) or decrease the number of living cells in the planktonic phase (a second antibiotic) decrease biofilm induction, while chemical inhibitors of cell lysis increase relative biofilm induction and shift the peak to higher antibiotic concentrations. Overall, our results offer experimental evidence linking cell wall synthesis inhibitors, cell lysis, increased eDNA, and biofilm formation in *E. faecalis* while also providing a predictive, quantitative model that sheds light on the interplay between cell lysis and antibiotic efficacy in developing biofilms.

## I. INTRODUCTION

Biofilms are dense, surface-associated microbial communities that play an important role in infectious diseases and a range of device-related clinical infections^1,2^. Biofilms exhibit a fascinating range of community behavior^3^, including long-range metabolic codependence^4^ and electrical signaling ^5–7^, phenotypic phase variation^8^ and spatial heterogeneity^9^, strong ecological competition^10^, and multiple types of cooperative behavior, including collective resistance to antimicrobial therapy^11–13^. The biofilm response to antibiotics has been a topic of particular interest, with biofilms across species showing dramatically increased resistance to antibiotics relative to planktonic cells. On the other hand, a number of studies have shown-somewhat counterintuitively-that exposure to sub-lethal doses of antibiotics may *enhance* biofilm formation in a wide range of species^14–16^. While antibiotic-mediated biofilm induction has been associated with modulated expression of biofilm-related genes, particularly those affiliated with bacterial and cell surface adhesion, cell motility, or metabolic stress, the mechanisms vary across species and drug classes and remain a focus of ongoing research efforts^15,16^.

In this work, we investigate the effects of sublethal antibiotic concentrations on biofilm formation in *E. faecalis*, a gram-positive bacteria commonly underlying nosocomial infections, including bacteremia, native and prosthetic valve endocarditis, and multiple device infections^1,17^. While our understanding of the molecular basis of both biofilm development and drug resistance in *E. faecalis* continues to rapidly mature^17,18^, surprisingly little attention has been paid to the impact of subinhibitory antibiotic treatments on *E. faecalis* communities. However, a recent series of intriguing studies has shown that *E. faecalis* biofilm formation (without antibiotic) hinges on an interplay between fratricide-associated cell lysis and the release of extracellular DNA (eDNA)^19–23^. More generally, eDNA is widely recognized as a critical component of biofilm structure in many species^24–26^. Additionally, a recent study in *S. aureus* showed that *β*-lactams administered at subinhibitory concentrations promoted biofilm formation and induced eDNA release in an autolysin-dependent manner^27^. Taken together, these results suggest that, for some drugs, biofilm induction hinges on a balance between the inhibitory effects of antibiotics-which reduce biofilm formation at sufficiently high concentrations–and the potential of antibiotic-induced cell lysis to promote biofilm formation, presumably through release of eDNA. Here we investigate this trade-off in *E. faecalis* biofilms exposed to multiple classes of antibiotics. We find that subinhibitory concentrations of cell wall synthesis inhibitors, but not other classes of drug, promote biofilm formation associated with increased cell lysis and increased eDNA and eRNA. Using a simple mathematical model, we quantify the trade-offs between drug efficacy and “beneficial” cell lysis and use the model to predict the effect of environmental perturbations, including the addition of DNase or chemical inhibitors of lysis, on the location and height of optimal biofilm production. Our results suggest that inhibitors of cell wall synthesis promote biofilm formation via increased cell lysis and offer a quantitative, predictive framework for understanding the trade-offs between drug toxicity and lysis-induced biofilm induction.

## II. RESULTS

### A. Cell wall synthesis inhibitors, but not other classes of antibiotics, promote biofilm formation at low concentrations

To investigate antibiotic induced biofilm formation, we exposed cultures of *E. faecalis* V583, a fully sequenced clinical isolate, to ampicillin during the first 24 hours of biofilm development. Using a bulk crystal violet staining assay (Methods), we observed a statistically significant enhancement of biofilm formation after 24 hours in the presence of low concentrations of ampicillin (Figure 1A). Ampicillin at these concentrations has almost no effect on growth of planktonic cultures, leading only to a slight decrease in the steady state cell density (Figure S1A). Similar enhancement of biofilm formation was observed for cells grown in different types of media (BHI, TSB) as well as for strain OG1RF, a common laboratory strain (Figure 1B), with the magnitude of the enhancement ranging from ≈ 10 – 30%.

**FIG. 1.**
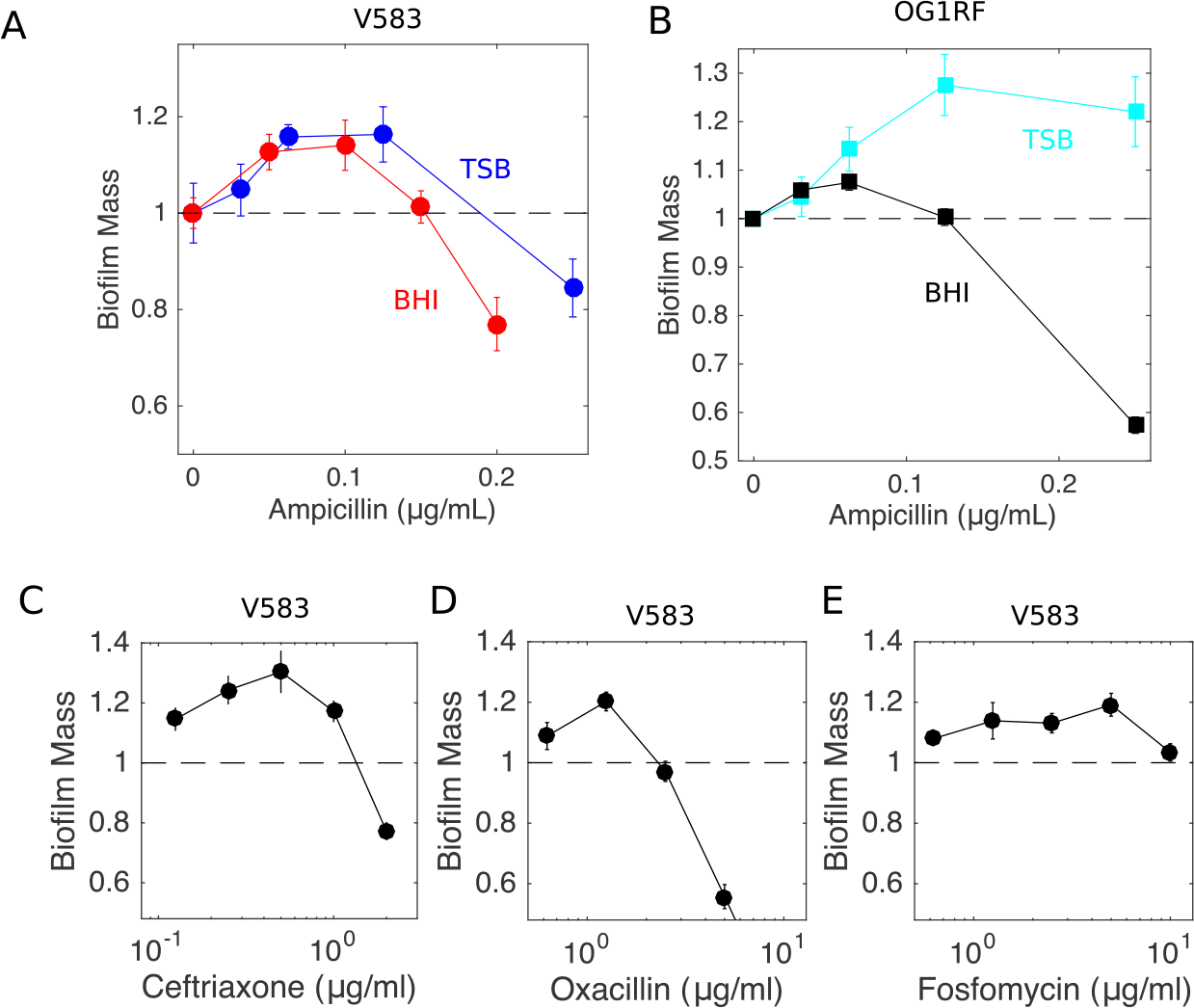
Inhibitors of cell wall synthesis enhance biofilm formation at low concentrations. A. Biofilm mass (normalized to 1 in the absence of drug) as a function of ampicillin concentration for E. faecalis strain V583 in TSB (blue) and BHI (red). B. Similar to panel A, with E. faecalis strain OG1RF in TSB (light blue) and BHI (black). Similar curves are also shown for V583 in BHI exposed to three additional cell wall synthesis inhibitors: ceftriaxone (C), oxacillin (D), and fosftomycin (E). In all panels, biofilm mass is measured by crystal violet assay (see Methods). Error bars are ± standard error of the mean from 6-12 replicates.

To determine whether the biofilm enhancement was specific to ampicillin, we performed similar experiments for antibiotics from multiple drug classes. Interestingly, we observed a similar increase of biofilm mass for other drugs inhibiting cell wall synthesis, including ceftriaxone, oxacillin, and fosfomycin (Figure 1C-E), whose mechanism of action is tightly linked to cell lysis. By contrast, drugs targeting protein synthesis, DNA synthesis, RNA synthesis, and folic acid synthesis did not appear to promote biofilm formation over the range of subinhibitory concentrations tested (Figure 2).

**FIG. 2.**
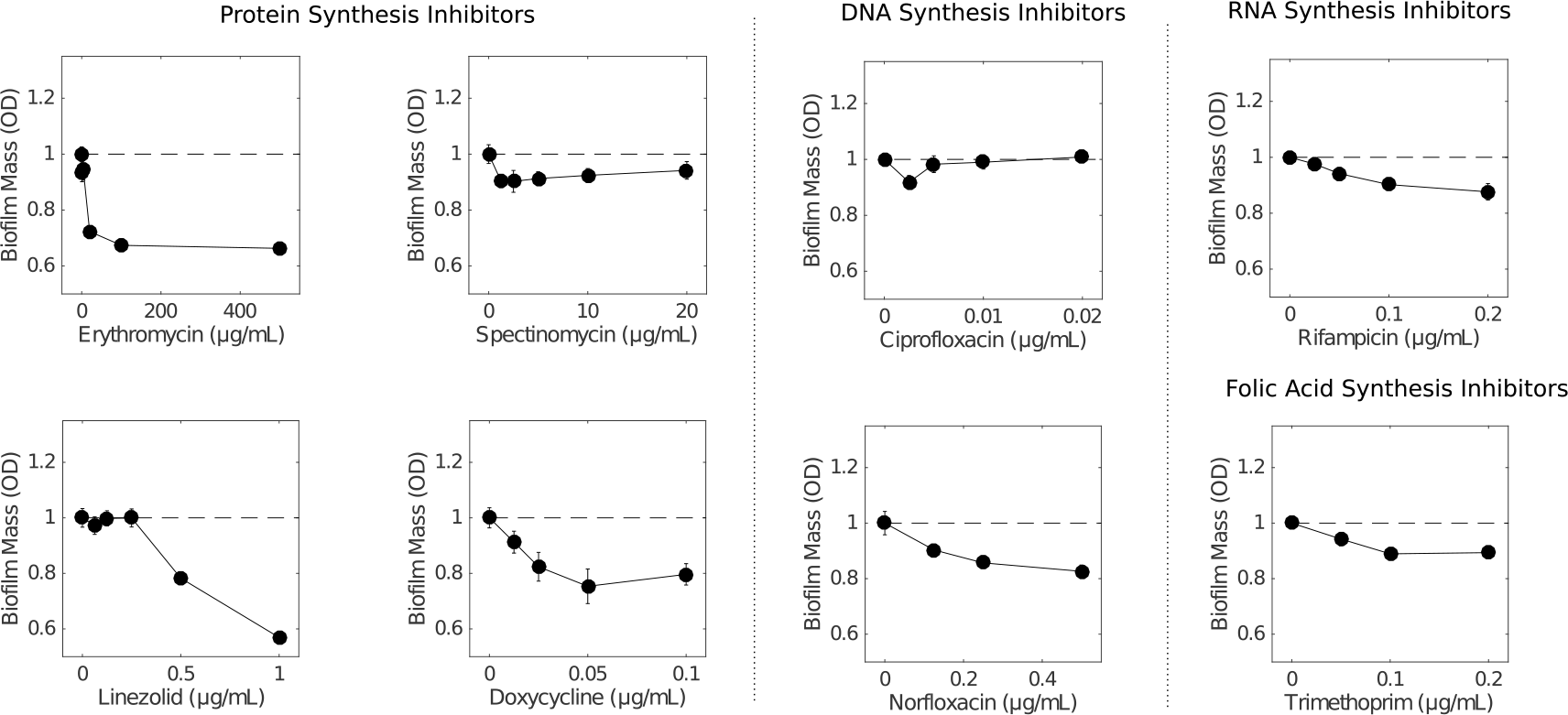
Antibiotics that do not target the cell wall do not enhance biofilm formation at low concentrations. Biofilm mass (normalized to 1 in the absence of drug) as a function of antibiotic for *E. faecalis* strain V583 in BHI exposed to protein synthesis inhibitors (erythromycin, spectinomcyin, linezolid, doxycycline), DNA synthesis inhibitors (ciprofloxacin, norfloxacin), RNA synthesis inhibitor (rifampicin), and folic acid synthesis inhibitors (trimethoprim). In all panels, biofilm mass is measured by crystal violet assay (see Methods). Error bars are ± standard error of the mean from 6-12 replicates.

### B. Biofilm enhancement occurs at subinhibitory concentrations but is associated with increased cell lysis and extracellular nucleic acid

In ampicillin, peak biofilm formation occurs for concentrations of approximately 0.1 *μ*g/mL, which has little effect on growth of planktonic cell cultures (Figure S1). To determine the effect of ampicillin on planktonic cultures over a wider drug range, we measured optical density time series of V583 cultures exposed to ampicillin concentrations up to 1 *μ*g/mL (Figure 3A). Ampicillin has little effect (< 15%) on the steady state density of cells up to concentrations of approximately 0.2 *μ*g/mL, and the dose response curve is well-approximated by a Hill-like function (commonly used in pharmacology^28^) with a half-maximal inhibitory concentration of *K*_50_ = 0.38 ± 0.01 *μ*g/mL. Therefore, increased biofilm formation occurs at concentrations considerably smaller than the half-maximal inhibitory concentration measured for planktonic cultures.

**FIG. 3.**
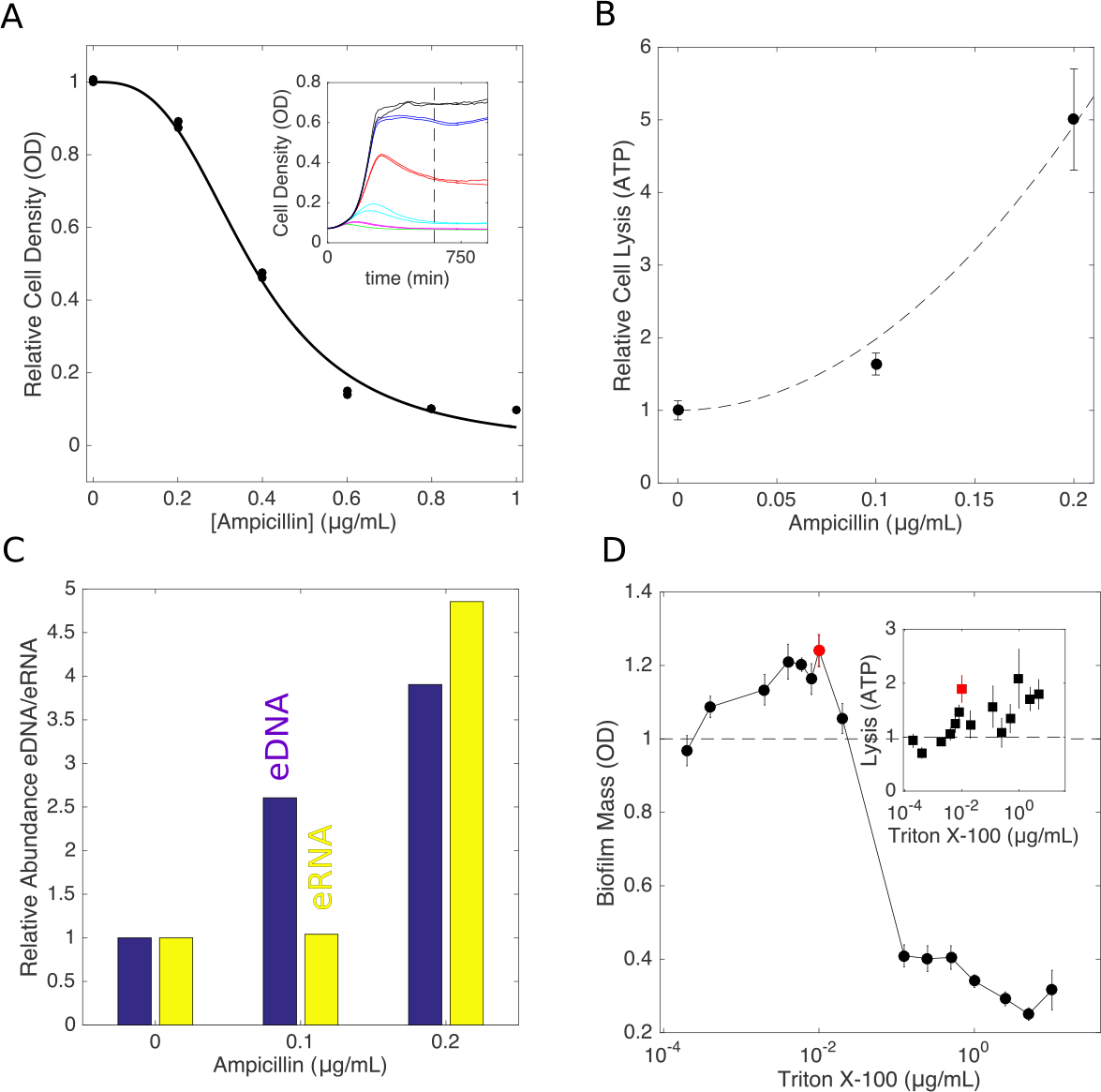
Enhanced biofilm formation occurs at sub-inhibitory concentrations and is associated with increased cell lysis and increased extracellular nucleic acid. A. Relative cell density (OD) approximately 10 hours after addition of ampicillin. Solid curve, fit to (1 − (*A*/*K*_50_)^*h*^)^−1^, with *A* the ampicillin concentration, *K*_50_ = 0.38 ± 0.01 *μ*g/mL the half maximal inhibitory concentration of the drug, and *h* = 3 a coefficient that describes the steepness of the dose response curve. Inset: time series of optical density following drug exposure at time *t* = 0 for ampicillin concentrations of 0 (black), 0.2 *μ*g/mL (blue), 0.4 *μ*g/mL (red), 0.6 *μ*g/mL (green), 0.8 *μ*g/mL (magenta), and 1.0 *μ*g/mL (cyan). B. Cell lysis (relative to untreated cells) as a function of ampicillin as measured by ATP luminescence assay (see Methods). Error bars are ± standard error of the mean from eight replicates. Dashed line, fit to 1 + *a*^2^/*r*_00_, with *a* the ampicillin concentration (measured in units of the drug’s half maximal inhibitory concentration (*K*_50_)) and *r*_00_ = 0.010 ± 0.001. C. Abundance of extracellular DNA (eDNA, blue) or RNA (eRNA, yellow) as a function of ampicillin concentration. Abundance is normalized relative to the eDNA (or eRNA) measured in the absence of drug. See also Figure S1. D. Triton X-100, a known inducer of cell lysis, enhances biofilm formation at low concentrations. Biofilm mass is measured by crystal violet assay (see Methods), and error bars are ± standard error of the mean from eight replicates. Inset: cell lysis (relative to untreated cells) as a function of Triton X-100 concentration. Red points correspond to peak in biofilm formation.

While these drug concentrations do not appreciably impact planktonic cell growth, it’s possible that they still produce a measurable increase in cell lysis. To investigate this issue, we measured cell lysis in 24 hour biofilms (Figure 3B) and planktonic cultures (Figure S1) using an ATP-based luminescence assay^29^. Indeed, we observed increased cell lysis even for low doses of ampicillin (≤ 0.2*μ*g/mL), with lysis increasing by nearly 5 fold in biofilms and several thousand fold in planktonic cultures for the highest doses.

Because eDNA has been implicated in *E. faecalis* biofilm formation, we next asked whether subinhibitory doses of ampicillin lead to increased quantities of extracellular nucleic acids in biofilms. To answer this question, we grew 24-hour biofilms in 5 mL cultures at various concentrations of ampicillin, harvested the biofilms and removed cells by centrifugation, and then extracted nucleic acid from remaining supernatant. We then quantified DNA (RNA) following treatment with RNase (DNase) using quantitative imaging of agarose gel electrophoresis (Figure 3C). Both eDNA and eRNA increase with ampicillin treatment, with eDNA (but not eRNA) increasing even at the lowest dose (ampicillin at 0.1 *μ*g/mL).

### C. Non-antibiotic induction of cell lysis promotes biofilm formation

Because cell lysis is observed at subinhibitory doses of ampicillin, and because lysis has been previously implicated in *E. faecalis* biofilm formation^19–23^, we next asked whether non-antibiotic inducers of cell lysis might also increase biofilm mass at small concentrations. To test this hypothesis, we grew biofilms in the presence of Triton X-100, a surfactant and known inducer of cell lysis^30^. Interestingly, we observed enhancement of biofilm formation similar in magnitude (≈ 20%) to that observed for cell wall inhibitors over Triton X-100 concentrations that yield similar (approximately 1.5-2 fold) increase of cell lysis (Figure 3D).

### D. Antibiotic-induced biofilm formation corresponds to an increase in the density of living cells and mean cell area

While our results indicate that biofilm mass is increased at low doses of ampicillin, it is not clear whether this enhancement is due to an increase in the number of living cells or merely an increase in bulk biofilm mass, which may include both viable and non-living components. To answer this question, we grew 16 replicate biofilms at 3 different antibiotic concentrations, treated them with live-dead cell stains, and quantified the number of live and dead cells in two-dimensional sections (i.e. the spatial cell density) at single-cell resolution using laser-scanning confocal microscopy (Methods). We observed an increase of approximately 25% in the number of living cells per slice, an increase similar in magnitude to the effects observed in bulk experiments (Figure 4). Interestingly, we also observed a slight increase in the mean size of a living cell as drug concentration increased. These results indicate that sections of biofilms formed under subinhibitory concentrations contain more living cells and more total living cell area–not merely an increase in non-living mass-than those formed in the absence of drug.

**FIG. 4.**
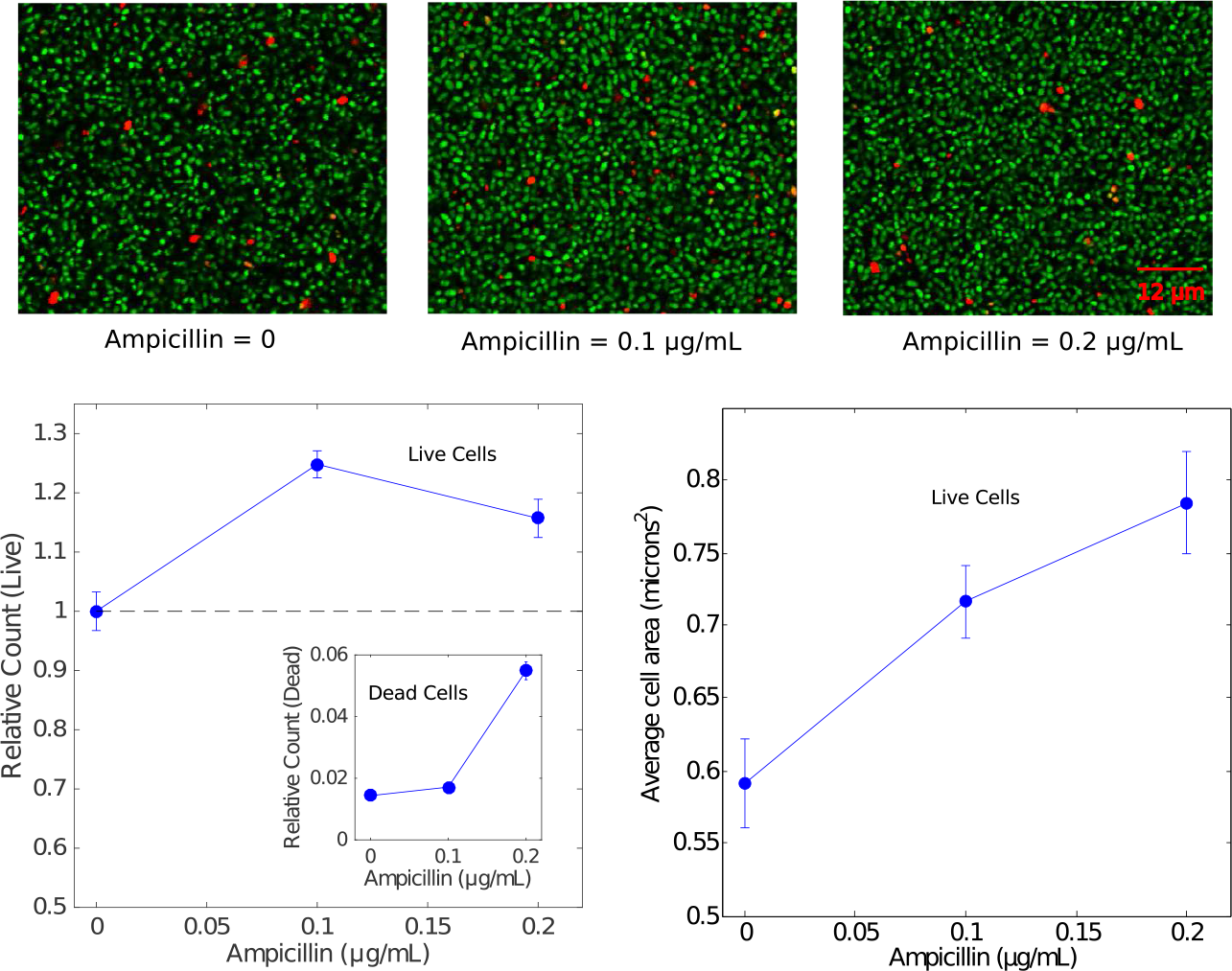
Enhanced biofilm formation corresponds to an increase in the density of living cells. Top panels: example sections from laser scanning confocal images of biofilms exposed to ampicillin at different concentrations (0, left panel; 0.1 *μ*g/mL, middle panel; 0.2 *μ*g/mL, right panel) and post-treated with live (green) and dead (red) stains. Lower left panel: Relative count of live cells and dead cells (inset) as a function of ampicillin concentration. Counts are normalized relative to the average number of live cells per 160 *μ*m × 160 *μ*m slice in the absence of drug, which is set to 1; note that each slice contains on the order of 10^3^ − 10^4^ live cells. Lower right panel: Mean area of living cells. Error bars are ± standard error of the mean taken over a total of 48 two dimensional slices per condition (three z-slices-taken at identical heights–for each biofilm and 16 independent biofilms per condition). The analysis involves on the order of 10^5^ total live cells per condition.

### E. A simple mathematical model describes biofilm induction as a balance between beneficial cell lysis and costly drug efficacy

To quantify the trade-offs between antibiotic efficacy and “beneficial” cell lysis, we developed a simple mathematical model describing the mass of living cells (*N*) and the mass of lysed cells and dead cell material (*D*), including eDNA, in a biofilm. Specifically, we have

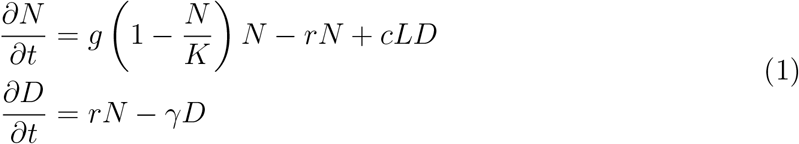

In the first equation, the first term describes logistic growth (with per capita growth *g* and carrying capacity *K* > 0), the second describes cell death (lysis) with rate *r* ≥ 0, and the last term describes the increase in biofilm mass due to surface attachment / adhesion of living cells in the planktonic phase (*L*), a process which is coupled to the dead cell mass *D* and controlled by a parameter *c* > 0. Models without this coupling do not exhibit a drug-induced maximum in biofilm mass (see SI). While the molecular mechanism of coupling is not specified in the model, this term could describe eDNA-mediated attachment and adhesion of planktonic cells, which is here assumed to occur at a rate proportional to both the living cells in solution (*L*) and the lysed cell material in the biofilm (*D*). In the second equation, the first term accounts for cell lysis and the second term describes a decay of lysed cell material (e.g. eDNA) due to, for example, detachment from the biofilm. The model implicitly assumes that the effect of antibiotic on cells in the planktonic phase occurs on a fast timescale, allowing *L* to reach a steady state on the timescale of biofilm formation. This assumption is consistent with experimental measurements, where planktonic populations reach a steady state size after approximately 10 hours (Figure 3), while biofilms are formed over a longer 24 hour timescale. The model includes two parameters, *r* = *r*(*a*) and *L* = *L*(*a*), that depend on drug concentration, *a*.

In the steady state, the living biofilm mass *N* is given by

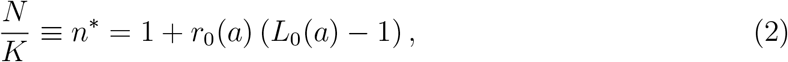

where *r*_0_(*a*) = *r*(*a*)/*g* and *L*_0_(*a*) = *cL*(*a*)/*γ* are (rescaled) functions describing the rate of cell lysis and the number of living cells in planktonic solution as a function of drug, *a*. Equation 2 illustrates a simple balance between the biofilm-inducing and biofilm-inhibiting effects antibiotics. To understand this tradeoff, we derived a phase diagram (Figure 5B) showing regions of enhanced biofilm formation-specifically, regions of the (*r*(*a*),*L*(*a*)) plane where *n*\* is greater in the presence of drug than in its absence (see SI for details). Enhanced biofilm formation is favored in regions of high lysis *r*(*a*) and large planktonic populations *L*(*a*). However, lysis *r*(*a*) is expected to increase with drug, while the number of available cells in planktonic phase *L*(*a*) is expected to decrease with *a* and eventually tend towards zero. The trade-off between these two effects determines the path taken by the system through the (*r*(*a*),*L*(*a*)) plane as drug is added. If increasing drug leads to increased lysis without a dramatic impact on the planktonic cells, the system exhibits enhanced biofilm formation (Figure 5B, path A). On the other hand, if increasing drug leads to a large decrease in planktonic cells and a relatively small increase in lysis, biofilm formation will not be enhanced (Figure 5B, path B).

**FIG. 5.**
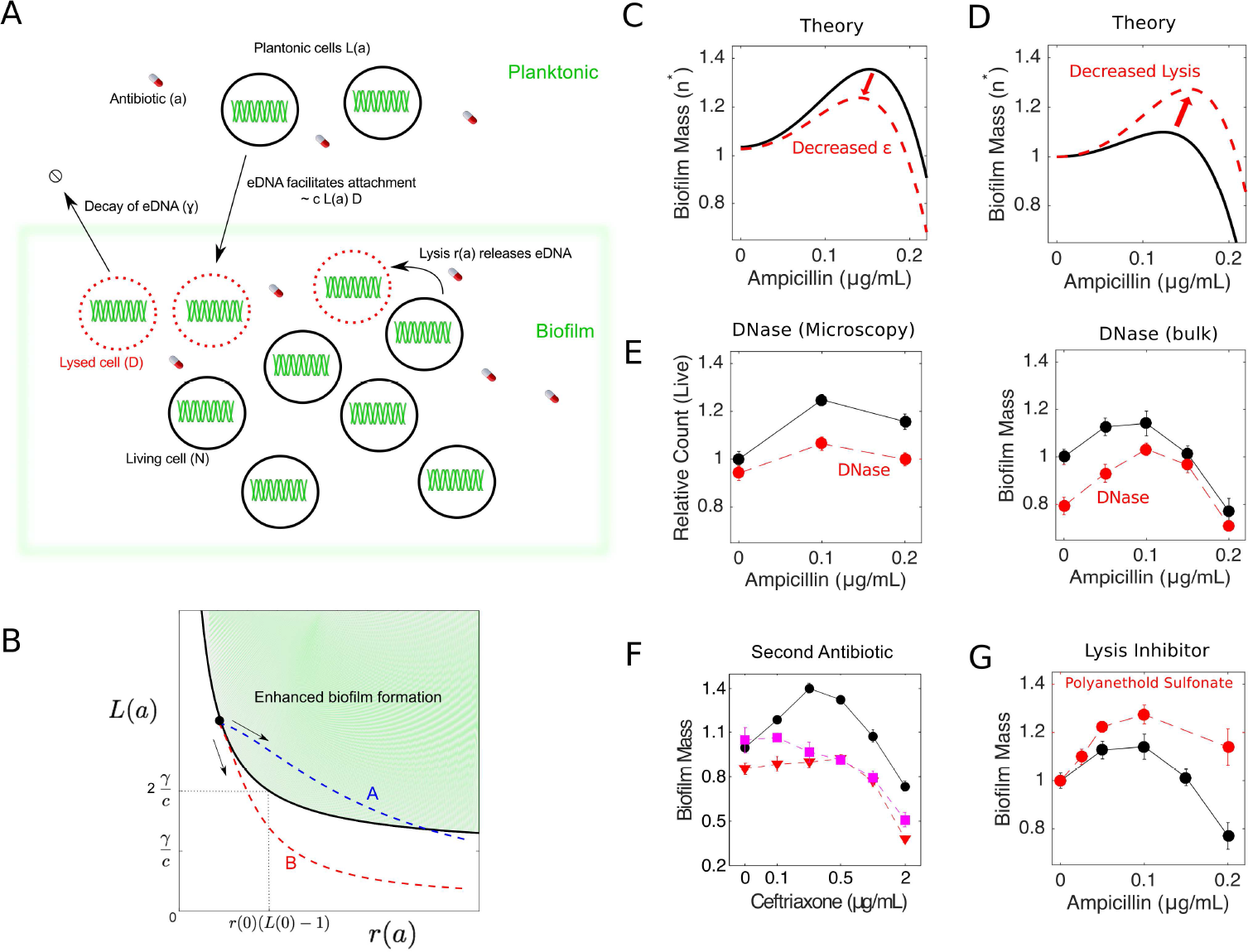
Mathematical model predicts effects of DNase, non lysis-inducing drugs, and cell lysis inhibitors. A. A simple mathematical model that couples cell lysis to biofilm formation describes qualitative features of antibiotic mediated biofilm enhancement. Lysis of living biofilm cells (*N*) depends on drug concentration *a* according to *r*(*a*). Lysed cells (*D*) facilitate cell adhesion and attachment from planktonic cells (*L*(*a*)). Adhesion/attachment is presumably enhanced due to release of eDNA, which itself detaches/decays at a rate *γ*, and surface attachment is proportional to *L*(*a*) and *D* with a rate constant *c*. In practice, the model contains two free “effective” parameters (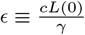 and 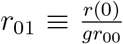) which can be estimated from the peak height and peak location in biofilm enhancement curves (e.g. Figure 1 or Figure 4). Intuitively, the parameter e describes the effective coupling between cell lysis and biofilm induction; a peak in biofilm mass occurs at nonzero drug concentration when *ϵ* > 1. B. Phase diagram shows the region of parameter space where enhanced biofilm formation occurs in terms of original model parameters. Enhanced biofilm formation is favored in regions of high lysis (*r*(*a*)) and large planktonic populations (*L*(*a*)). Blue (red) dashed lines: path taken by system as antibiotic concentration is increased; arrows indicate direction of increasing *a*. The blue curve (path A) exhibits enhanced biofilm formation, while the red curve (path B) does not. C. The model predicts that perturbations that decrease *ϵ* (left panel) lead to a decrease in the height and location of the peak in living biofilm mass (*n*\* ≡ *N*/*K*, solid line). Parameter values *ϵ* = 1.18 ± 0.01 and *r*_01_ = 19 ± 4 in the absence of perturbation are estimated from living biofilm cell counts from confocal microscopy (Figure 4). Dashed curve shows the predicted change in peak location and height due to perturbations that reduce the coupling e by several percent. D. The model predicts that perturbations that decrease cell lysis lead to an increase in the height and location of maximum living biofilm mass (*n*\*, solid line). Parameter values *ϵ* = 1.09 ± 0.02 and *r*_01_ = 18 ± 6 are estimated from bulk experiments (Figure 1). Dashed curve shows the predicted change in peak location and height due to perturbations that decrease cell lysis. E. Relative biofilm mass (solid curves) as a function of ampicillin from confocal microscopy (left; see also Figure 4) and bulk experiments (right; see also Figure 1). Dashed curves: identical experiments but with DNase I added at a concentration of 0.4 mg/mL. F. Relative biofilm mass as a function of ceftriaxone alone (solid curve, circles) or ceftriaxone in combination with a constant concentration of rifampicin at 0.3 *μ*g/mL (squares, dashed) or tetracycline at 0.2 *μ*g/mL (triangles, dashed). G. Solid curve: same as in panel E, right. Dashed curves: identical experiments but with sodium polyanethole sulfonate (SPS), a known inhibitor of cell lysis, at a concentration of 10 *μ*g/mL. Error bars represent ± standard error of the mean.

In principle, the functional forms for *r*_0_(*a*) and *L*_0_(*a*) could be derived from microscopic models that describe the molecular level dynamics of antibiotic-induced cell lysis and cell death. Fortunately, however, the functions *r*_0_(*a*) and *L*_0_(*a*) can also each be estimated–up to a scaling constant–by independent experimental measurements, even in the absence of a detailed molecular model. Specifically, using the data in Figure 3A-B, we take

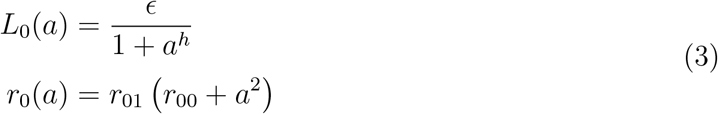

where *h* = 3 describes the steepness of the dose response curve (and is analogous to Hill coefficients used in biochemistry to describe cooperative binding between ligands), *a* is measured in units of the drug’s half-maximal inhibitory concentration, estimated to be *K*_50_ = 0.38 ± 0.01 *μ*g/mL, and *r*_00_ = 0.010 ± 0.001. It should be noted that we assume a quadratic dependence of lysis on *a* to match the experimental measurements; this should be viewed as a simple parameterization of the experimental lysis measurements and does not imply any particular mechanism. The quadratic dependence of lysis on *a* could depend on complex pharmacological and pharmacodynamics of the antibiotics, and we do not attempt to model those here.

The remaining two parameters, *ϵ* and *r*_01_, are scaling parameters that can be estimated from biofilm induction curves (e.g. Figure 1). Because the measured value of *r*_00_ << 1, we can derive approximate solutions for the location (*a*_*max*_) and height (*p*_*h*_) of the biofilm peak using simple perturbation theory (SI). Specifically, the peak location is given by

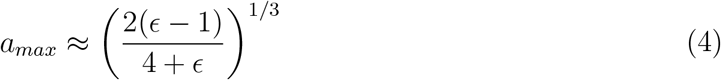

and the peak height is given by

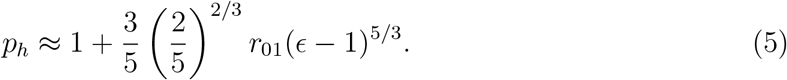

These expressions indicate that an optimum in biofilm production occurs at a nonzero concentration *a* when *ϵ* > 1, and the effect of further increasing *ϵ* is to shift the peak to higher *a* and increase its height. We refer to *ϵ* as an effective coupling parameter, and given the functional forms in Equation 3, the value of *ϵ* alone determines whether there is a peak in biofilm formation at nonzero drug concentrations. In terms of the original model parameters, *ϵ* is given by 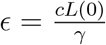, where *L*(0) is the size of the planktonic cell population at zero drug concentration, and *r*_01_ is given by 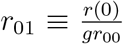, where *r*(0) is the native level of lysis in the absence of drug. Antibiotic-mediated biofilm formation occurs for *ϵ* > 1 and is therefore favored by large rates of lysis-mediated adhesion (*c*), high concentrations of planktonic cells (*L*(0)), and slow degradation / decay of eDNA (*γ*).

### F. Model predicts effects of DNase treatment, a second antibiotic, and lysis inhibitors

While it is straightforward to estimate *ϵ* and *r*_01_ from biofilm experiments–for example, *ϵ* = 1.09 ± 0.02 and *r*_01_ = 18 ± 6 based on the bulk experiments in Figure 1A–it is more instructive to consider the qualitative predictions of the model as parameters are varied. Our model predicts that perturbations that decrease *ϵ* will lower the peak height (Figure 5B, left panel; Figure S2, bottom left). On the other hand, perturbations that decrease cell lysis would shift *r*_0_(*a*) → *r*_0_(*a*) − *β*;, with *β* a positive constant (or equivalently, decreasing lysis would shift *r*_00_ → *r*_00_ − *β*/*r*_01_). While the latter effects would not be evident at the level of the approximate equations (Equations 4, 5), we can evaluate the predicted effects numerically (Figure 5C, left panel, and Figure S2, bottom right panel) or by looking at higher-order terms in the approximation (SI). Decreasing lysis is predicted to shift the peak location to higher drug concentrations and, somewhat counterintuitively, leads to an increase in biofilm induction (that is, an increase in the height of the peak relative to the drug-free case). In words, a higher concentration of antibiotic is needed to achieve sufficient cell lysis to induce increased biofilm production.

To test these predictions experimentally, we first repeated both bulk and microscopy experiments in the presence of DNase I. Because eDNA has been implicated as the molecular conduit linking cell lysis to biofilm formation, we expect DNase treatment to decrease *ϵ* by effectively increasing the decay rate *γ*. Indeed, biofilms treated with DNase exhibit lower peaks (Figure 5D). The model also predicts a slight shift in the location of the peak, but the resolution of the experimental data is insufficient to evaluate that prediction quantitatively. A second way of decreasing *ϵ* would be to decrease the number of living cells in planktonic phase (α). One possibility is to treat the cells with a second (non-lysis-inducing) antibiotic, which is expected to lower the steady state density of cells in liquid phase. Consistent with this prediction, we observed that treatments with a cell wall synthesis inhibitor (ceftriaxone) along with a second, non-lysis inducing drug (tetracycline or rifampicin) decrease the height of the peak to almost zero (Figure 5E).

Next, to test the prediction that decreasing lysis leads to an increase in relative peak height and location, we repeated the biofilm experiment in the presence of sodium polyanethole sulfonate (SPS). SPS is a common anticoagulant used in clinical blood cultures and is known to have a positive effect on bacterial survival^31^. It has been shown to inhibit of cell lysis in staphylococci by suppressing activity of autolytic wall systems^27,32,33^. To verify that SPS inhibits cells lysis in our system, we measured lysis as a function of SPS concentration using an ATP-based luminescence assay^29^.; our data suggests that SPS inhibits cell lysis by approximately 40% in the absence of drug at the concentrations used (Figure S3). We found that treatment with the lysis inhibitor appears to shift the peak to slightly higher drug concentrations and increases the magnitude of biofilm enhancement, again consistent with predictions from the model.

## III. DISCUSSION

Our work demonstrates that biofilm formation in *E. faecalis* is enhanced by subinhibitory concentrations of cell-wall synthesis inhibitors, but not by inhibitors of protein, DNA, folic acid, or RNA synthesis. Enhanced biofilm formation is associated with increased cell lysis and an increase in eDNA and eRNA. We observed similar enhancement effects when cultures were treated with non-antibiotic chemicals that induce similar amounts of cell lysis. To quantify the trade-off between drug toxicity and the beneficial effects of cell lysis, we developed a simple mathematical model that predicts changes to drug-induced biofilm formation due to external perturbations that reduce eDNA, reduce living cells in the planktonic phase, or inhibit cell lysis. Our model suggests that antibiotic-induced biofilm formation occurs when the drug-induced increase in cell lysis is sufficiently large relative to the drug-induced decrease of living cells in the planktonic phase.

Subinhibitory concentrations of antibiotics have been reported to promote biofilm formation in multiple species via a range of different mechanisms^15,16^. However, relatively little is known about drug-induced biofilm formation in *E. faecalis*. Subinhibitory antibiotic concentrations have previously been shown to impact the physioelectrical^34^ and adhesion behavior^35^ of *E. faecalis*. In addition, low concentrations of tigecycline have been shown to reduce biofilm formation, even when growth of planktonic cells is not significantly affected^36^. To our knowledge, however, this is the first work to describe enhancement of biofilm formation due to cell wall synthesis inhibitors in *E. faecalis*. On the other hand, our work suggests that inhibitors of protein, DNA, folic acid, and RNA synthesis do not promote biofilm formation over a wide range of subinhibitory concentrations (though we do caution that we cannot definitively rule out biofilm promotion at higher concentrations, perhaps via different mechanisms).

Our results are consistent with the established role of eDNA in biofilm formation and may be applicable to drug-induced biofilm formation in other species, most notably *S. aureus*^27^. However, it is possible that other mechanisms may also contribute–at least in part–to our results. For example, recent work has shown that eDNA is prevalent in biofilms even at the early developmental stages when cell lysis is minimal ^29^, and it is possible that subinhibitory drug concentrations increase eDNA through similar non-lysis mechanisms. In addition, it is well-known that sub-MIC levels of antibiotic can dramatically alter gene expression profiles in bacteria^37–39^, indicating that biofilm enhancement may arise from a complex combination of multiple factors. Finally, we note that live-dead cell staining results should be interpreted with some caution, because uptake of various stains may be variable^40–45^. Nevertheless, our results are promising because they suggest that, at least in the experimental regimes measured here, a simple conceptual (and mathematical) model is sufficient to describe and predict the primary effects of drug exposure.

Despite the model’s success, it is without question a dramatic oversimplification of the complex biofilm formation process. Computational models of biofilm formation are a powerful tool for understanding dynamics and evolution of complex communities^46–48^, and detailed models may contain dozens or even hundreds of microscopic parameters. Yet even the most elaborate mathematical models neglect biological details at some scale. Our approach was not to develop a detailed microscopic model, but rather to develop a simple, minimal model to help intuitively explain and predict the trade-offs between antibiotic efficacy and beneficial cell lysis at the population level. Linking our model with more detailed agent-based simulations may help us further understand the potential role of spatial structure and heterogeneity in drug-induced biofilm formation. For example, recent work has shown that in the absence of drug, *E. faecalis* biofilm formation depends on a phenotypic bistability in gene expression, giving rise to lysis-susceptible and lysis-inducing sub-populations^19–23^. It would be interesting to further explore the interplay between this multi-modal population structure and drug-induced lysis observed in this work.

Our work also raises intriguing questions about how genetic resistance determinants might spread in biofilm populations, even in the absence of the strong selection pressure of high drug concentrations. A quantitative understanding of biofilm formation may also inspire new optimized dosing protocols, similar to those in, for example,^49–51^, and the current model could be easily extended to investigate the effects of clinically realistic antibiotic dosing regimens. In the long run, these results may lay the groundwork for improved, systematic design of biofilm-specific therapies^52,53^.

## IV. METHODS

### A. Bacterial strains and media

Experiments were performed with *Enterococcus faecalis* V583, a fully sequenced clinical isolate^54^, and strain OG1RF, which was derived from human oral isolate OG1 ^55^. For each experiment, starting cultures (3 mL) were inoculated from a single colony on brain heart infusion (BHI) agar plates (1.5% (w/v) bacteriological agar) and grown overnight in BHI or tryptic soy broth (TSB) medium at 37°C without shaking.

### B. Drugs

Antibiotics used in this study are listed in table I. Antibiotic stock solutions were sterilized by passing through 0.22 *μ*m filter, aliquoted into daily-use volumes, and kept at −20 or −80°C for no more than 3-6 months. All chemicals and media were purchased from Sigma-Aldrich or Fisher Scientific unless stated otherwise.

**TABLE I.** Antibiotics used in this study

| Antibiotics | Description | Abbreviation |
| --- | --- | --- |
| Ampicillin | Cell wall synthesis inhibitor | AMP |
| Ceftriaxone | Cell wall synthesis inhibitor | CRO |
| Fosfomycin | Cell wall synthesis inhibitor | FOF |
| Oxacillin | Cell wall synthesis inhibitor | OXA |
| Doxycycline | Inhibit 30s subunit | DOX |
| Spectinomycin | Inhibit 30s subunit | SPT |
| Erythromycin | Inhibit 50s subunit | ERM |
| Linezolid | Inhibit 50s subunit | LZD |
| Ciprofloxacin | DNA synthesis inhibitor | CIP |
| Norfloxacin | DNA synthesis inhibitor | NOR |
| Rifampin | RNA synthesis inhibitor | RIF |
| Trimethoprim | Folic Acid synthesis inhibitor | TMP |

### C. Growth curves of *Enterococcus faecalis*

Overnight cultures were diluted 100X into fresh BHI medium, and then 200 *μ*L of diluted culture were added to each well of a 96-well clear bottom plate. Different concentrations of antibiotics were then added to each well, and time series of optical density (OD600) were measured at 15 minute intervals for 24 hours at 30°C using an EnSpire Multimode Plate Reader in a temperature controlled warm room.

### D. Microtiter plate biofilm assay

We measured biofilm mass in bulk assays using a well-established crystal violet staining assay^56,57^. Overnight cultures were diluted 100X into fresh BHI medium, and then 100 *μ*L of diluted culture (along with appropriate concentrations of antibiotics, if relevant) were added to each well of a flat-bottomed polystyrene microtiter 96-well plate (Greiner Bio-One Cellstar). The plate was incubated at 37°C without shaking for 24 hours. After incubation, supernatant from liquid cultures was removed by gently turning over the plate, shaking, and patting on paper towels. Wells were then gently washed with PBS. To fix the biofilm on the plate, 125 *μ*L 96% ethanol was added into each well and allowed to incubate for 20 minutes. Ethanol was then removed and the plate was dried at room temperature for half an hour. Following drying, 125 *μ*L 0.5% crystal violet was then added to stain the biofilm mass. After 30 minutes, plates were washed multiple times with fresh PBS. Plates was then turned upside down and dried for 1 hour. Finally, 125 *μ*L 30% acetic acid was added to each well in order to dissolve biofilm. Solutions were transferred to a new 96-well plate and absorbance readings at 590 nm were taken using Enspire multimodal plate reader. For each treatment, we performed 6-12 replicates.

### E. ATP detection assay

To measure cell lysis, we used the luminescence assay described in^29^ to measure increases in extracellular ATP using a commercial ATP Determination kit (Molecular Probes). Prior to measuring cell lysis, biofilms were grown as previously described with different concentrations of antibiotics in a 96-well plate for 24 hours. We then washed the plate twice with nuclease-free water and removed excess liquid. After washing, 10 *μ*L nuclease-free water was added to each well and biofilms were scraped down by using inoculation loops or pipette tips. Solutions were transferred to a new 96-well white polystyrene plate (Thermo Scientific Nunc F96 MicroWell) and 90 *μ*L ATP standard assay solution from ATP Determination kit (Molecular Probes) was added to each sample. Luminescence was measured by plate reader.

### F. Confocal laser scanning microscopy

Bacterial cultures (200 *μ*L total volume, with appropriate drug treatment) were grown in replicates of 4 in 16-well chambered coverglass vessels. After incubating for 24 hours, liquid was removed and plate was washed twice with filtered millipore water and then stained using LIVE/DEAD BacLight Bacterial Viability kit (Molecular Probes) for 20 minutes. After staining, liquid was removed and coverglass was afixed to the top of the chamber.

The biofilms were imaged using a Zeiss LSM700 confocal laser scanning microscope (40X, 1.4 N.A. objective (Zeiss)) with laser lines 488 nm and 555 nm used for exitation. For each well, four image stacks (160 × 160 microns) spanning 20-30 microns (vertically) at 1 micron intervals were taken at four separate (x,y) locations on the cover slip, giving a total of 16 biofilm images per condition. To analyze images, we split them into red and green channels, set the threshold for each slice individually using automated thresholding algorithms in ImageJ, and then used a watershed algorithm to segment cells and determine location and size of each cell type (live / dead) in each slice. Cell counts per slice were averaged over three well-separated slices (to avoid double counting cells in adjacent slices) in the middle portion of each biofilm and over all 16 images per condition.

### G. Extracellular DNA/RNA extraction

Biofilms were grown in triplicate with ampicillin 0, 0.1 and 0.2 *μ*g/mL in 6-well polystyrene plates with a total volume of 5 mL for each well. After 24 hours, we dumped out liquid and washed the plate twice with PBS before adding 1 mL 1X Tris-EDTA (TE, 10mM Tris-Cl, 1 mM EDTA, pH=8.0) buffer and scraping biofilms from bottom of plates.

After harvesting biofilms, cells were removed by centrifugation and supernatant was purified by using only the binding and washing steps in QIAprep Spin Miniprep kit according to the manufacturer’s instruction. 5 volumes of PB buffer was added to 1 volume of supernatant and mixed. 800 *μ*L of solution was transferred to a spin column and centrifuged at 13000 rpm for 1 minute. A volume of 0.5 mL PB buffer was added to wash the spin column followed by centrifugation for 1 minute. Then 0.75 mL PE buffer was added to spin column, and the column was centrifuged again for 1 minute. The flow-through was discarded and residual was removed by centrifuging spin column for an additional 1 minute. The column was then transferred to a new 1.5 mL microcentrifuge tube and 30 *μ*L EB buffer was added to the center of the spin column. After 1 minute, the column was centrifuged for 1 minute to elute DNA. DNase I or RNase was added to the same treatment samples as controls.

### H. Agarose gel electrophoresis

Gel tray and all related tools were rinsed with nuclease-free water, and samples were loaded on a 1% agarose gel with 1X Tris-acetate-EDTA (TAE, pH=8.4) buffer containing an appropriate volume of SYBR safe. The gel was run at 120V for 40 mins in 1X TAE buffer. DNA or RNA fragments were virtualized under UV light from UV transilluminator. To analyze images, ImageJ software was used to subtract background and perform intensity analysis for different lanes. Identically sized regions were selected for different lanes and a profile plot of each lane was drawn. A straight line across the base of the peak was drawn to enclose the peak, and the wand tool was used to select each peak and measure percentage of relative densities.

